# Determinant host differences in ZIKV sfRNA accumulation

**DOI:** 10.1101/2025.11.17.688941

**Authors:** Taissa Ricciardi-Jorge, Skye Storrie, Daniel Santos Mansur, Trevor R. Sweeney

## Abstract

Zika virus (*Orthoflavivirus* zikaese) is part of the *Orthoflavivirus* genus that contains several viral species posing major and expanding burdens to public health worldwide. Upon infection, orthoflaviviruses produce viral RNA fragments named sfRNAs (short flaviviral RNA), generated from viral genomic RNA incompletely degraded by Xrn1/Pcm, the major cellular 5’-3’ exonuclease. Several conserved elements in the orthoflavivirus 3’ untranslated region (UTR) are responsible for <u>X</u>rnI-resistance, named <u>xr</u>RNA. The ubiquitous prevalence of xrRNA/sfRNA in flaviviruses indicate an important role of these structures in the viral life cycle, although their function is not completely understood. Here, we used *in vitro* reconstitution and infection models to examine the role of ZIKV xrRNA2 in mammalian and insect cells. SHAPE RNA structure probing revealed that disruption of ZIKV xrRNA2 does not affect xrRNA1 or dumbbell structure elsewhere in the 3’UTR despite disrupting all sfRNA accumulation in mammalian cells. *In vitro* RNA degradation assays showed that xrRNA1 efficiently stalled human or mosquito Xrn1 independent of xrRNA2 integrity. Reversion in mammalian cells occurred at early passages in protein kinase R (PKR)^-/-^ or IFNα-receptor1 (IFAR1)^-/-^/PKR^-/-^ knock out (KO) or double KO mammalian A549 cells, respectively, while wild-type virus outcompeted the xrRNA2 defective virus in RNAi-deficient mosquito cells. Together, our data reveals novel details of the mechanism of sfRNA accumulation in mammalian and insect cells We demonstrate a differential sensitivity to the absence of a functional xrRNA2 structure in mammalian and insect cells, with reversion to an active xrRNA2 occurring more rapidly in mammalian cells. Together, our work reveals new details on the requirements for ZIKV sfRNA accumulation in mammalian and mosquito cells.

## INTRODUCTION

The *Orthoflavivirus* genus contains several viral species that represent major and expanding burdens in public health worldwide, including Zika virus (ZIKV), dengue virus (DENV) and West Nile virus (WNV), among others (1, 2). Click or tap here to enter text. Orthoflavivirus genomes consist of a linear positive-sense single-stranded RNA ∼11kb in length that contains a long open reading frame flanked by highly structured 5’ and 3’ untranslated regions (UTRs) (3-5). The secondary structure of orthoflavivirus 3’UTRs contains several conserved and often duplicated elements shared among viruses that circulate between vertebrate and invertebrate hosts, as well as invertebrate-exclusive viruses (5) Click or tap here to enter text.. Some of these conserved elements can resist Xrn1 5’-3’ degradation, hence being referred to as “XrnI-resistant RNAs” (xrRNA). As a result, during infection, 3’ fragments of incompletely degraded genomic RNA termed short flaviviral RNA fragments (sfRNAs) are also produced during infection alongside the full-length genomic RNA (6-10).

The number and length of sfRNAs vary between different viruses but both their wide prevalence and duplication and redundancy (11) indicates an important role of these RNAs with various functions reported during infection. The sfRNAs in ZIKV and other orthoflaviviruses have been reported to enhance virus replication by various mechanisms. For example, DENV2 sfRNA sequesters TRIM25 to inhibit interferon (IFN) expression (12) and G3BP1, G3BP2 and CAPRIN1 to inhibits IFN-simulated gene (ISG) translation (13), ZIKV sfRNA was reported to stabilise expression of the viral RdRp, NS5, enhancing inhibition of IFN signalling (14). Interestingly, the impact of sfRNA production can vary between the mammalian and insect vector host. ZIKV sfRNA production was required for dissemination into the foetal brain in a mouse model of infection and caused apoptosis of infected neural progenitor cells (14). SfRNAs also enhanced viral dissemination in mosquitoes but lead to a decrease in apoptosis in mosquito tissues (15). Given their apparent importance for enhancing viral replication, the role and formation of sfRNAs is increasingly studied as a potential mechanism for attenuated virus vaccine design.

The atomic structures of different xrRNAs reveal complex tertiary interactions in which the 5’ end threads a ring-like structure of RNA formed by stabilising interactions, including pseudoknots (PK), and base stacking, that vary between different viruses (9, 16, 17). The secondary structures of ZIKV xrRNA1 and -2 are shown in Figure 1B with key contacts important for tertiary folding highlighted. Additional dumbbell (DB) structures can also arrest Xrn1 within the viral 3’UTR resulting in sfRNAs of varying length accumulating within a single infection (18, 19). Accumulation of different sfRNAs during infection highlights that Xrn1 can penetrate xrRNAs under certain conditions that are not currently understood. Interestingly, mutation of DENV or ZIKV xrRNA2 disrupted all sfRNA generation in mammalian cells (19, 20) with the authors suggesting that different xrRNAs may interfere with others function. Separately, disruption of the DB structure in the ZIKV 3’UTR also disrupted accumulation of sfRNA production from all xrRNAs (18).

**Figure 1.**
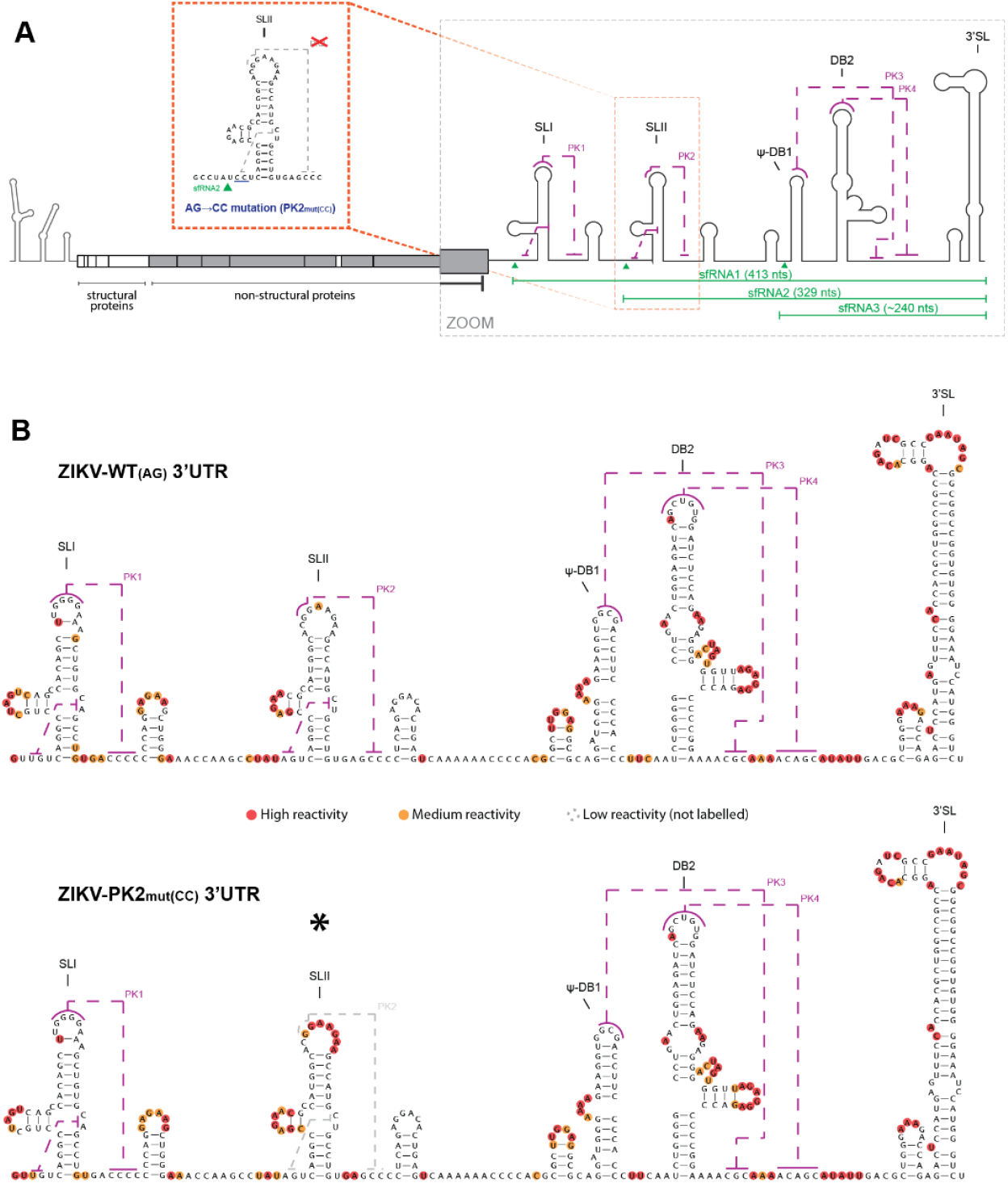
Two-nucleotide mutation confirmed by structural mapping to disrupt only pseudoknot 2 formation. (**A**) Schematic representation of ZIKV genome with zoomed details of 3’UTR secondary structure and local of AG→CC mutation designed to disrupt PK2/xrRNA2. (**B**) Results of nucleotide reactivity profile by SHAPE of ZIKV-WT_(AG)_ and -PK2_mut(CC)_ 3’UTR.

Here, we used *in vitro* reconstitution and infection models to examine the role of ZIKV xrRNA2 in mammalian and insect cells. We use selective 2’-<u>H</u>ydroxyl <u>A</u>cylation analysed by <u>P</u>rimer <u>E</u>xtension (SHAPE) RNA structure probing to show that disruption of ZIKV xrRNA2 does not affect xrRNA1 or DB structure elsewhere in the 3’UTR despite disrupting all sfRNA accumulation in mammalian cells. *In vitro* RNA degradation assays showed that xrRNA1 efficiently stalled human or mosquito Xrn1 independent of xrRNA2 integrity. Reversion in mammalian cells occurred at early passages in protein kinase R (PKR)^-/-^ or IFNα-receptor1 (IFAR1)^-/-^/PKR^-/-^ knock out (KO) or double KO mammalian A549 cells, respectively, while wild-type virus outcompeted the xrRNA2 defective virus in RNAi-deficient mosquito cells. Together, our data reveals novel details of the mechanism of sfRNA accumulation in mammalian and insect cells We demonstrate a differential sensitivity to the absence of a functional xrRNA2 structure in mammalian and insect cells, with reversion to an active xrRNA2 occurring more rapidly in mammalian cells. Together, our work reveals new details on the requirements for ZIKV sfRNA accumulation in mammalian and mosquito cells.

## RESULTS

### Structural changes caused by PK2 disruption are localised

Numerous base pairing contacts are required to maintain the integrity of RNA secondary and tertiary structures (Figure 1). To specifically disrupt sfRNA2 formation with minimal interference to the overall UTR structure, we aimed to destabilize PK2, responsible for Xrn1 resistance at this location of the viral genome (xrRNA2) (6, 8). To do so, mutations on the two key nucleotides (AG→CC) immediately preceding SL2 were introduced to the viral genome (PK2_mut(CC)_), shown in Figure 1A. To confirm the desired alteration in the RNA tertiary structure, we examined the ZIKV 3’UTR, by using <u>S</u>elective 2’-<u>H</u>ydroxyl <u>A</u>cylation analysed by <u>P</u>rimer <u>E</u>xtension (SHAPE) RNA structure analysis, as previously described (21)Click or tap here to enter text..

SHAPE results present high reactivity in loop regions, while base-paired regions are less reactive. Exceptions to this trend typically occur where loops are predicted to form tertiary structures, such as sites of pseudoknot formation. Nucleotide reactivities, mapped on the wildtype (WT_(AG)_) 3’UTR structure (Figure 1B), were comparable to previously published SHAPE results and structural predictions (22, 23). In contrast to previous studies, we obtained SHAPE reactivity data for the entire 3’UTR by incorporating an ordered 3’ extension to accommodate the primer binding site for the reverse transcription reaction.

SHAPE reactivities for the PK2_mut(CC)_ 3’UTR were highly consistent with those of the WT_(AG)_ RNA except for the region around the SL2/PK2/xrRNA2 (marked in the figure with an asterisk). The increased nucleotide reactivity at the loop of SL2 confirmed the that the AG→CC mutation disrupted the PK2 formation without alteration of the SL2 secondary structure. The reactivities in the remainder of UTR were unchanged overall, indicating an intact secondary and tertiary structure elsewhere, including the xrRNA1 structure for which no structural changes were detected. Hence, we confirmed that the mutation used in this study disrupts only the tertiary structure responsible for PK2, as intended.

### Single nucleotide reversion emerged in mammalian cells restores PK2/xrRNA2 and sfRNA formation

Using a well-characterized ZIKV reverse genetic system (24) Click or tap here to enter text.the mutation AG→CC was incorporated into the virus genome. ZIKV-WT_(AG)_ and -PK2_mut(CC)_ were recovered following transfection of *in vitro* transcribed and capped viral genomic RNA into Vero cells (Figure 2A), a cell line chosen for its impaired type 1 IFN production (25). The recovered virus (P0) was titrated and its sequence verified before being used for further viral characterization. The PK2_mut(CC)_ virus was also passaged five times in Vero cells for testing mutation stability, which was revealed to be low, leading to single nucleotide reversion, CC→CG (PK2_rev(CG)_), detected by Sanger sequencing as soon as passage 2 (Figure 2B). A mock and a ZIKV-WT_AG_ were parallelly passaged five times in Vero cells as controls. No emerging mutation was detected in the 3’UTR and no cross-contamination were detected in the mock.

**Figure 2.**
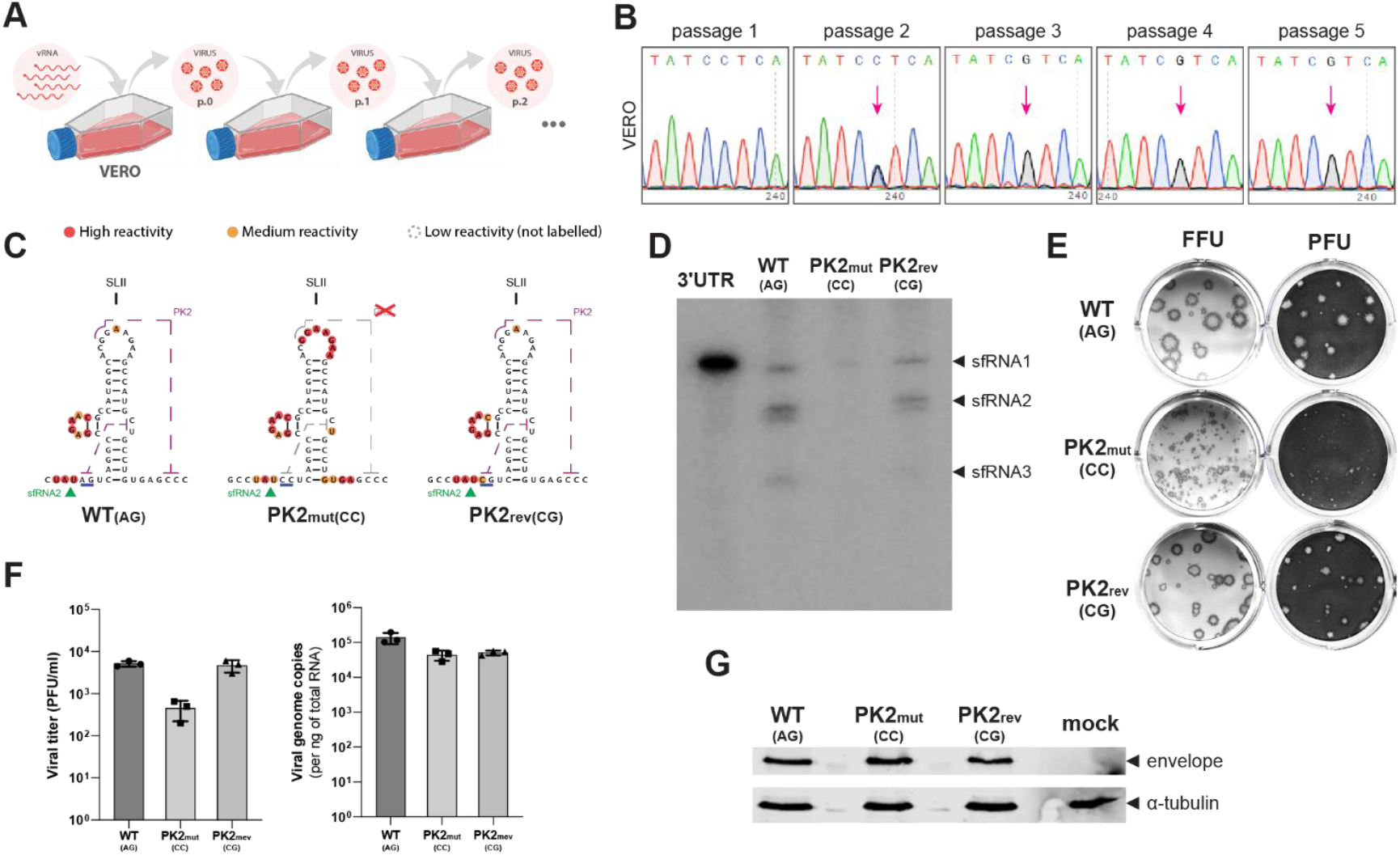
PK2mut(CC) virus disrupt total sfRNA synthesis is under negative selective pressure in mammalian cells despite so significant fitness loss. (**A**) Schematic representation of viral passage numbering of virus rescue from reverse genetics in Vero cells and consecutive viral passages. (**B**) Electropherogram of Sanger sequencing results obtained during five serial passages of ZIKV-PK2_mut(CC)_ in Vero cells, focused on mutation site. Emerging C→G reversion is highlighted by a pink arrow. (**C**) SHAPE results on SL2 region of 3’UTR of ZIKV-WT_(AG)_, -PK2_mut(CC)_, and -PK2_rev(CG)_. (**D**) Northern blot of Vero cells infected with ZIKV-WT_(AG)_, -PK2_mut(CC)_, and -PK2_rev(CG)_ for 48 hours. Resolved in denaturing gel and probed for ZIKV-3’UTR. (**E**) Plaque and foci phenotype of ZIKV-WT_(AG)_, -PK2_mut(CC)_, and -PK2_rev(CG)_ in Vero cells after staining with toluidine blue (upper wells, PFU) or immunostaining with anti-ZIKV envelope protein and secondary antibody conjugated to alkaline phosphatase and reacted with chromogenic substrate (lower wells, FFU). (F-G) Vero cells were infected with ZIKV-WT_(AG)_, -PK2_mut(CC)_, and -PK2_rev(CG)_ at MOI 1 and incubated for 24 hours. (**F**) Viral titre in infection supernatant quantified by PFU/ml and viral genome copies analysed by RT-qPCR from cell lysates. (**G**) Cell lysates were also analysed by western blot, immunostaining with antibodies directed against Flavivirus E protein and alpha-tubulin.**Selective pressure to keep ZIKV PK2/xrRNA2 structure in mammalian cells is not linked to ISR or IFN pathways**

By investigation with SHAPE, it was possible to verify the loss of reactivity on the loop of SL2 of PK2_rev(CG)_ comparable to WT_AG_ virus, which shows that the single-nucleotide reversion CG was enough to restore the PK2 structure (Figure 2C). Further investigation by northern blot revealed that the disruption of PK2/xrRNA2 abrogated all sfRNA formation in Vero cells infected with the PK2_mut(CC)_ virus (Figure 2D). Upon recovery of tertiary structure on the PK2_rev(CG)_ virus, the pattern of sfRNA formation was restored to levels similar to the WT_(AG)_ virus.

Phenotypical characterization of these viruses shows that the plaque formation in Vero cells was greatly affected in the PK2_mut(CC)_ virus and mostly recovered on the PK2_rev(CG)_ virus (Figure 2E). Despite that, analysis of viral protein production, genome copies and viral titres in Vero cells (Figure 2F-G) showed no significant differences between the WT_(AG)_, PK2_mut(CC)_ or PK2_rev(CG)_ viruses.

Altogether these results show that, despite the absence of measurable effect of the virus replication, the disruption of PK2/xrRNA2, and the abrogation of sfRNA, in the PK2_mut(CC)_ virus is under strong negative selective pressure.

Human A549 immortalised cells derived from lung epithelia display a robust IFN driven antiviral response in comparison to Vero cells(26, 27). When examined in A549 cells, the PK2_mut(CC)_ virus did not demonstrate any differences in viral replication parameters when compared to WT_(AG)_ virus (Figure 3A). As in Vero cells, the passage of this mutant virus in A549 cells led to the same single-nucleotide reversion (CC→CG) detectable by Sanger sequencing as soon as passage 2 (Figure 3B).

**Figure 3.**
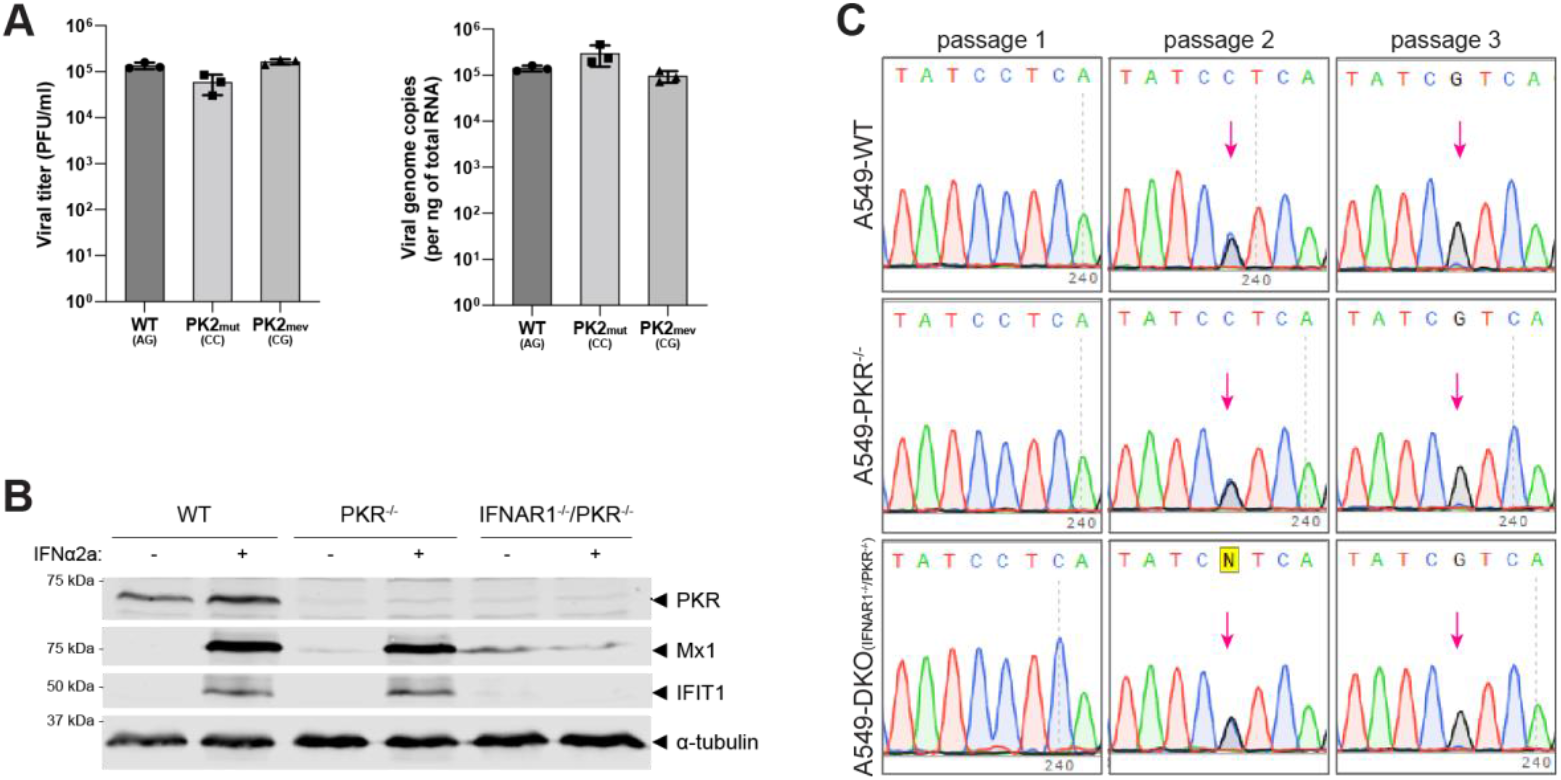
Investigation of selective pressure on ZIKV-PK2_rev(CG)_ during passages in mammalian cell lines. (**A**) A549 wildtype cells were infected with ZIKV-WT_(AG)_, -PK2_mut(CC)_, and -PK2_rev(CG)_ at MOI 1 and incubated for 24 hours. Viral titre in infection supernatant quantified by PFU/ml and viral genome copies analysed by RT-qPCR from cell lysates. (**B**) Western blot characterizing A549 cell lineages (WT, PKR^-/-^ and IFNAR1^-/-^/PKR^-/-^) used for viral passages, with or without stimulation with 100U/ml of IFNα2a for 24 hours. (**C**) Electropherogram of Sanger sequencing results obtained during five serial passages of ZIKV-PK2_mut(CC)_ in A549-wildtype (upper panels), -PKR^-/-^ (middle panels) and -PKR^-/-^/IFNAR1^-/-^ (lower panels) cells, focused on mutation site. Emerging C→G reversion highlighted by pink arrow. Sequencing results presented are representative of three independent passage experiments.

Previous studies have associated the sfRNA function with the inhibition of IFN-I signalling (14, 20) and PKR activation (28) in mammalian cells. However, viral passages of PK2_mut(CC)_ virus performed in PKR-knockout and PKR/IFNAR1-double knockout cells led to the emergence of the same single-nucleotide reversion (CC→CG) at passage 2, as the passages in VERO and wildtype A549 cells (Figure 3B). These results show that the PKR-dependent ISR and IFN pathways have no impact on the selective pressure observed in mammalian cells to restore PK2/xrRNA2 structures and sfRNA production. The consistency of the selective pressure over the virus, weakens the hypothesis that the observed selective pressure to restore the integrity of PK2/xrRNA2 may be due to its consequences over total sfRNA synthesis.

### Selective pressure to keep ZIKV PK2/xrRNA2 structure in insect cells is not linked to RNAi pathway or total sfRNA abrogation

The stability of the PK2_mut(CC)_ virus was next tested by serial passages in insect cell lines, that are known to offer less selective pressure over flavivirus evolution than mammalian cells (5). C6/36, a RNAi-deficient cell line (29) was chosen to explore previous association of flavivirus sfRNA with RNAi pathway in insect cells (30-32). Interestingly, the same single-nucleotide reversion (CC→CG) observed in mammalian cells was detectable at passage 4 in the insect cell lines (Figure 4A), showing that a consonant evolutive pressure to restore PK2 is observed in the cells of both vertebrate and invertebrate hosts. The decreased viral fitness of the PK2_mut(CC)_ virus compared to the WT_AG_ virus was confirmed by competition assay. Equal titres of WT_AG_ and PK2_mut(CC)_ were used to infect C6/36 cells but after two passages, the WT_(AG)_ virus was dominant (Figure 4B).

**Figure 4.**
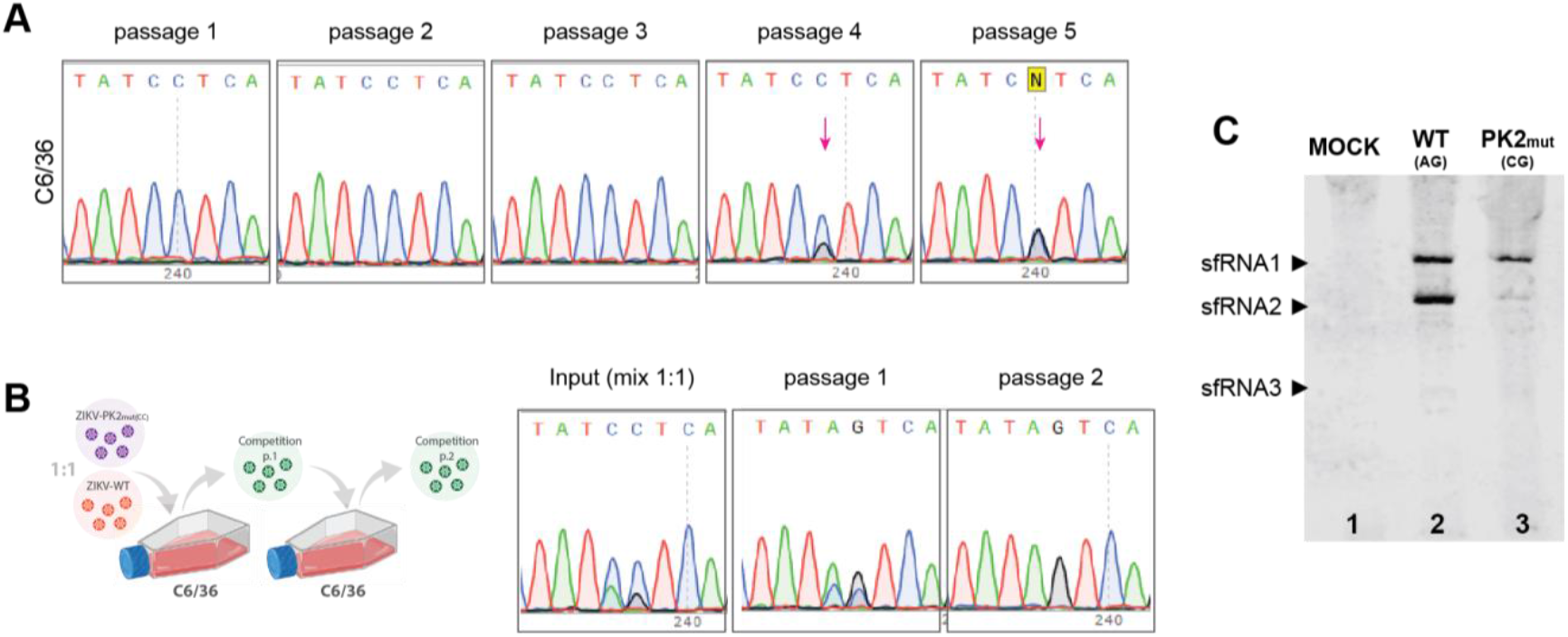
Investigation of ZIKV-PK2_rev(CG)_ stability in insect cell lines. (**A**) Electropherogram of Sanger sequencing results obtained during five serial passages of ZIKV-PK2_mut(CC)_ in C6/36-wildtype cells, focused on mutation site. Emerging C→G reversion highlighted by pink arrow. (**B**) Schematic representation and electropherogram of Sanger sequencing results of competition assay of ZIKV-WT_(AT)_ and ZIKV-PK2_mut(CC)_ carried out in C6/36 cells, starting from 1:1 mix of both viruses and passaged twice. (**C**) Northern blot of C6/36 cells infected with of ZIKV-WT_(AG)_, -PK2_mut(CC)_, and -PK2_rev(CG)_ for 72 hours. Samples volumes to an equivalent of 10^10^ ZIKV genome copies/slot resolved in denaturing gel and probed for ZIKV-3’UTR. Northern blot and sequencing results presented are representative of three independent experiments.

Further investigation by Northern blot has shown that, in accordance with previously published results (15, 20) in insect cells, the disruption of PK2/xrRNA2 only disturbs the formation of sfRNA2 (Figure 4C). Altogether, these results weaken the hypothesis that the strong negative pressure on PK2/xrRNA2 disruptions observed in mammalian cells is due to the abrogation of total sfRNA. Instead, the consistency of the selective pressure over the virus, despite the cell type suggests that the integrity of PK2/xrRNA2 may be instead a requirement of the virus itself and, therefore, host independent.

### PK2/xrRNA2 disruption does not affect xrRNA1 function in sfRNA1 formation by Xrn1 in *in vitro* digestion assay

Inline with other reports (14, 15) we observed that infection of mammalian cells with PK2_mut(CC)_ virus reduced both sfRNA1 and sfRNA2 (Figure 2D), while infection of insect cells, only disrupted sfRNA2 (Figure 4G), as expected by SHAPE mapping showing that that PK2_mut(CC)_ mutation left xrRNA1 unaffected (Figure 1). To test if this difference could be due to species specific differences in enzymatic activity or processivity of Xrn1, the catalytic domains of Xrn1 from *H. sapiens* and the orthologue Pcm from *A. aegypti* were expressed in *E*.*coli* (Figure 5A). These enzymes, together with commercially available Xrn1 from *S. cerevisiae*, were tested in an *in vitro* digestion of fragments of the WT_AG_ and PK2_mut(CC)_ genome.

**Figure 5.**
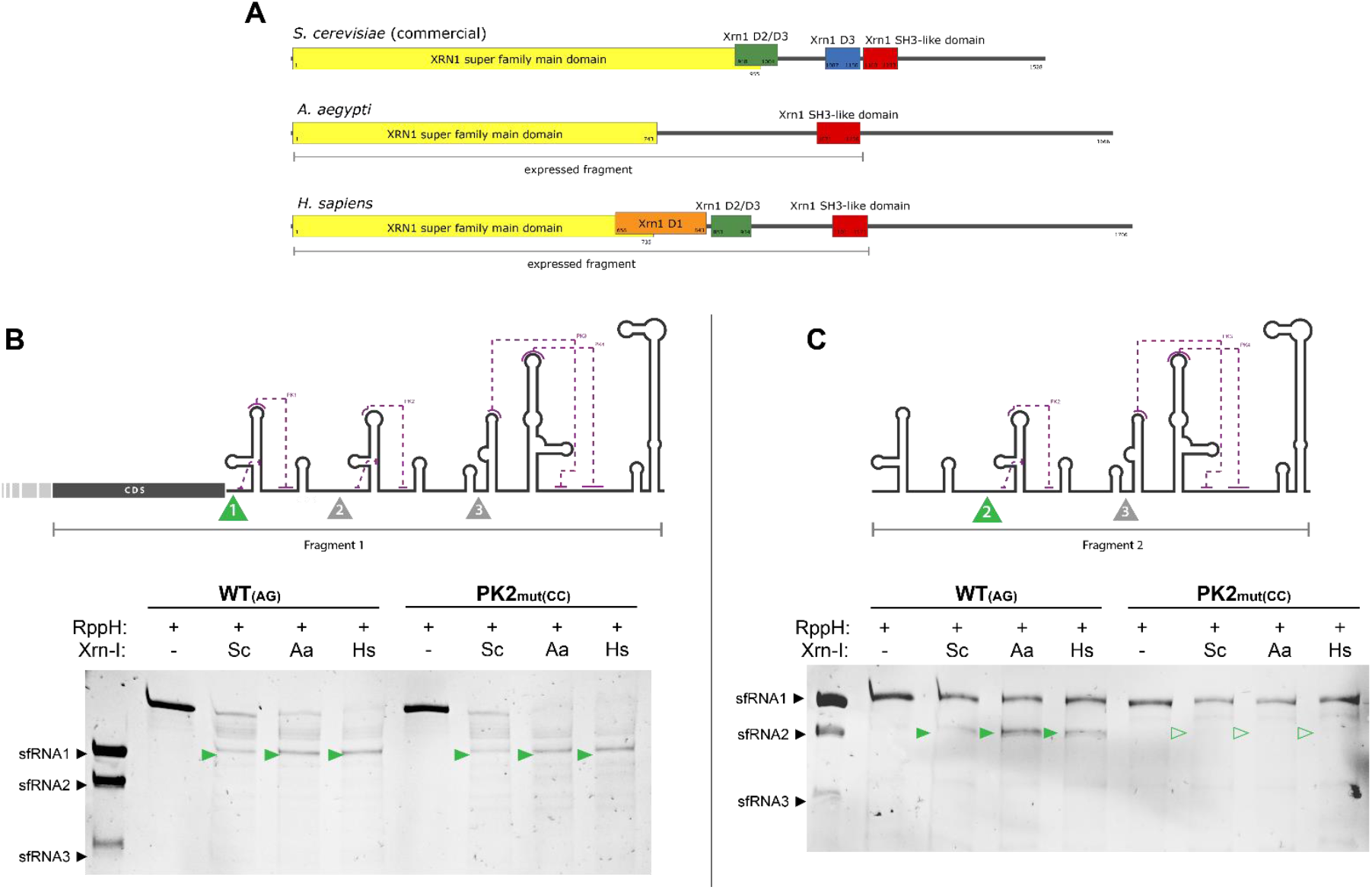
Xrn1 *in vitro* digestion of 3’UTR fragments of ZIKV-WT_(AG)_ and -PK2_mut(CC)_. (**A**) Schematic representation of Xrn1 proteins from *S. cerevisiae, H. sapiens* and *A. aegypti* and respective domains. (**B**) Schematic representation of fragment designed to test pseudoknot 1 integrity and its ability to halt Xrn1 exonuclease activity generating fragments equivalent to sfRNA1, signed by a green arrow with number 1. Representative result of *in vitro* digestion assay resolved in acrylamide gel under denaturing conditions. The full green arrows highlight the fragment corresponding to sfRNA1. (**C**) Schematic representation of fragment designed to test pseudoknot 2 integrity and its ability to halt Xrn1 exonuclease activity, generating fragments equivalent to sfRNA2, signed by a green arrow with number 2. Representative result of *in vitro* digestion assay resolved in acrylamide gel under denaturing conditions. The full green arrows highlight the presence of RNA fragments corresponding to sfRNA2 and the hollow arrows its absence. The results shown are representative of three independent experiments.

As observed in Figure 5, xrRNA1 was equally effective at halting Xrn1-degradation whether or not the downstream xrRNA2 was disrupted (Figure 5B). In contrast, while PK2/xrRNA2 from WT_AG_ virus halted exonuclease degradation, the modified PK2/xrRNA2 from PK2_mut(CC)_ virus did not (Figure 5C). No difference was observed when comparing Xrn1 processivity between vertebrate and invertebrate hosts. Therefore, while our data confirm a role for xrRNA2 in promoting accumulation of all ZIKV sfRNAs in mammalian but not insect hosts, we demonstrate that this is not due to an impact on structure or function of other RNA elements in the viral 3’ UTR, or difference in processivity of exoribonucleases from different species.

## DISCUSSION

In this work we designed a two-nucleotides mutation (AG→CC) adjacent to SL2, named PK2_mutCC_, to disrupt only ZIKV PK2/xrRNA2 without affecting SL2 secondary structure or any secondary/tertiary structure on the rest of the UTR, which we confirmed using RNA SHAPE structure analysis. An alternative mutational strategy to disrupt xrRNA activity is to mutate a conserved cytosine residue important for xrRNA stability (14, 15). Interestingly, mutation of both xrRNAs in ZIKV using this method was reported to abolish virus recovery in mammalian or insect cells; reversion of one site to restore either sfRNA 1 or sfRNA2 reproducibly occurred (14). In contrast, complete deletion of both xrRNA-1 and -2 structures in ZIKV was tolerated and the double deletion virus was recovered from insect cells (15, 20) providing a possible route to attenuated ZIKV vaccine development. The reason behind the striking difference in recovery of mutated versus deleted xrRNA viruses is not known but highlights the importance of further study on xrRNA function and sfRNA accumulation for basic and translational research.

The disruption of PK2/xrRNA2 showed alteration of plaque phenotype but not of viral fitness in mammalian cells. The alteration of plaque phenotype upon disturbances of sfRNA synthesis has been previously reported for ZIKV (18) and other orthoflaviviruses (33). The little impact of viral fitness upon mutation on PK2/xrRNA2 is consistent with previous studies of ZIKV (15, 18) Click or tap here to enter text.Despite that, the mutation must have a cost for the virus, as PK2_mutCC_ showed low stability upon serial passages in mammalian cells (Vero and A549), presenting the emergence of a single nucleotide reversion CC→CG in mammalian cells as early as in passage 2. These results agree with previous studies that have observed the strong requirement of PK2 presence/integrity for adaptation in mammalian cells (5, 20). PK2_mutCC_ was also shown to lead to major disruption of sfRNA synthesis in these cells, not only of the corresponding sfRNA2, a phenomenon previously observed in ZIKV (14), DENV (19) and JEV (33), but curiously not in WNV (6, 7).

The single-nucleotide reversion CC→CG was shown to be enough to restore the PK2/xrRNA2, and by consequence sfRNA formation, which could suggest that the selective pressure for PK2 restoration was due to the lack of sfRNA, as hypothesized by previous studies (20). The presence of sfRNA would be important for virus replication as it would play a role in counteracting cellular antiviral responses via IFN-signalling (14) and by ISR, that in ZIKV-infected A549 is driven only by PKR (26, 28). However, viral passages in PKR^-/-^ and IFNAR1^-/-^/PKR^-/-^ cells lead to the same negative selection of the PK2_mutCC_ virus observed in the wildtype cells, indicating that this phenomenon is IFN- and ISR-independent. These results agree with the absence of impact over viral fitness observed for the PK2_mutCC_ virus. If the mutation could impact in such important antiviral pathways, its consequences over virus replication would be very significant, which is, indeed, not observed.

Upon passage of PK2_mutCC_ virus in insect cells, known to offer less selective pressure over flavivirus evolution than mammalian cells (5, 34, 35) we observed the emergence of the same CC→CG reversion previously observed. The reduced viral fitness of PK2_mutCC_ virus was confirmed in this model by competition assay. This is in contrast with results obtained with DENV2, that showed high tolerance for mutations and deletions around SL2 upon passages in insect cells and in mosquitoes (5). The reversion CC→CG of PK2_mutCC_ virus was observed in mosquito cells deficient in RNAi (29), indicating that this reversion is independent of a reported role for sfRNA in evading the RNAi response (30-32), but more consistent with alternative studies of sfRNA function (15, 36).

The remarkable consistency of reversion observed in this study, dispute the hypothesis that the strong negative pressure on PK2/xrRNA2 disruptions observed in mammalian cells is due to the abrogation of total sfRNA in these cells. It also suggests that the function of PK2 is not made redundant by PK1, as per seen in insect cells. Instead, this consistent reversion implies that the role for sfRNA accumulation and/or intact 3’UTR structural elements may extend beyond host innate immune evasion mechanisms and promotes a viral fitness advantage either in a host-independent manner, via interaction with viral proteins or other portions of the viral RNA; or by some conserved protein/pathway in both host cells. Consistent with this idea, Cerikan et al. identified a surprisingly critical role for the DENV 3’UTR in establishing virus replication organelles indistinguishable from those observed during infection (37). ZIKV sfRNA has also been recently reported to affect autophagy pathways (37) and also to sequester the ZIKV polymerase NS5 in the cytoplasm (14). As NS5 translocation to the nucleus was reported to impact cellular gene expression in DENV-2 infection (38) the possibility of further disruption of cellular function by increased nuclear accumulation of NS5 in the absence of sfRNA cannot be excluded. Further studies are therefore necessary to investigate the subtle but consistent viral fitness advantage given by an intact PK2.

Finally, the PK2 disruption revealed a difference in ZIKV sfRNA2 formation in mammalian and insect cells. In insect cells, PK2_mutCC_ virus only presents disruption of sfRNA2 synthesis, and not also of upstream sfRNA1 as seen in mammalian cells, in accordance with other studies in insect cells (15, 20) but the reason for this has not been investigated. One possible explanation for the differential processing of the same RNA sequence could be from the temperature in which each cell line is maintained, as the interactions that stabilizes PKs are dynamic (7). Alternatively, differences in Xrn1 sequences between hosts-human and mosquito Xrn1 share 45% identity-could alter sfRNA accumulation. Our results of Xrn1 *in vitro* digestion assay however contest both possibilities, as PK1/xrRNA1 of WT_AG_ and PK2_mutCC_ virus could equally halt activity of yeast, mosquito and human Xrn1 at 37°C. This result is in accordance with the observation that, unlike in mosquito-derived cell lines, when whole mosquitoes are infected by PK2-mutated/deleted virus there is a disruption of sfRNA synthesis as observed in mammalian cells (6, 23). Therefore, the difference in sfRNA accumulation in wild-type PK2 and PK2mut CC’s in mammalian and insect cells is potentially due to altered interactions with host proteins, which requires further investigation. Previous structure modelling of sfRNAs from DENV, WNV and ZIKV based on small angle X-ray scattering experiments suggested that sfRNAs have extended structures potentially accommodating individual domains (22) with no apparent interaction between xrRNA-2 and DB regions. It is important to note that these modelling experiments were performed on isolated sfRNAs. Given the propensity of sfRNA to bind to viral and host proteins, it is possible that binding of a mammalian factor helps stabilise ZIKV sfRNAs during infection and that an intact xrRNA-2 (this study) and DB structure (18) is required for this interaction. Conversely, these intact structures may interfere with a mammalian factor(s) that binds and targets sfRNA for degradation. Together, our findings develop our understanding of sfRNA accumulation during infection, a critical host/pathogen interface in this important family of pathogenic viruses.

## MATERIAL AND METHODS

### Cells and viruses

Vero and A549 cells were grown at 37°C/5% CO2 in DMEM media supplemented with 10% foetal bovine serum. PKR^−/−^ and DKO (IFNAR^−/−^/PKR^−/−^) cell line has been previously described (26). C6/36 cells (ATCC CRL-1660), Aag2 cells were grown at 28°C in L-15 medium supplemented with 10% foetal bovine serum and tryptose (2 g/litre). Aag2-AF5 cells (ECACC 19022601), a clonal population of Aag2 cell line, and Aag2-AF319 (ECACC 19022602), a clonal population containing Dcr2 knockout mutation (Dcr2-^/-^), were used in this study.

ZIKV was recovered from previously described reverse genetics system (24). Viral RNA was transcribed *in vitro* with in house-produced enzyme and capped with ScriptCap™ m7G Capping System (CellScript, cat. no. C-SCCE0625). The capped RNA was electroporated into Vero cells using Neon™ Transfection System (Invitrogen) following protocol provided by the fabricant. The virus recovered from electroporated cells at the onset of CPE was considered passage 0 (P0). The mutations of interest (AG→CC for PK2_mut(CC)_, and AG→CG for PK2_rev(CG)_) were introduced to the reverse genetic plasmid by PCR (CC forward: AAGCCTAT<u>CC</u>TCAGGCCGAGAACG; CC reverse: TCTCGGCCTGA<u>GG</u>ATAGGCTTGG; CG forward: AAGCCTAT<u>CG</u>TCAGGCCGAGAACG; CG reverse: TCTCGGCCTGA<u>CG</u>ATAGGCTTGG). All viral titrations were performed in Vero22 cells by plaque-forming unit (PFU) or focus-forming unit (FFU).

### Viral infections and passages

All viral infection were performed in media without foetal bovine serum supplementation for 90 min. After virus absorption the cells were washed in PBS and then incubated in complete media. Viral passages were performed in independent triplicates having an uninfected control in parallel, always after infection at MOI 0.001. Passages in mammalian cells were incubated for 48 hours, while passages in insect cells were incubated for 3 days. From the infection, cells were collected for RNA extraction and sequencing after reverse transcription step; and infection supernatant was collected in aliquots for viral titration and subsequent infection.

### SHAPE analysis

For ZIKV 3’UTR analysis by SHAPE, a DNA fragment corresponding to the end of ZIKV genome was produced by PCR (forward primer: <u>TAATACGACTCACTATA</u>GGCCTTCGGGCCAAGCACCAATCTTAATGTTGTCAGG; reverse primer: <u>GAACCGGACCGAAGCCCGATTTATAGTAAATGGATCCGGCGAACCGGATCT</u>AGACCCA TGGATTTCCCCAC) from the reverse genetic plasmid of interest. The amplicon generated includes a T7 promoter and extends the 3’ end to include a double stem-loop structure that folds independently of the 3’UTR RNA structures. The double stem-loop structure provides an alternative primer binding site to the ZIKV 3’UTR 3’SL to ensure SHAPE reactivity data is obtained for the entire 3’UTR in in a single reaction at optimal coverage. This amplicon was transcribed *in vitro* with in house-produced enzyme and, following purification step, the RNA was submitted to SHAPE according to previously published protocols (21, 39, 40). The NMIA-modified RNA and control were used as template for reverse transcription using primer (GAACCGGACCGAAGCCCG) labelled with VIC (NMIA-modified RNA) or NED labelled (unmodified RNA used as ladder, reverse transcribed in the presence of ddCTP). Reaction products were analysed by capillary electrophoresis and results by QuShape, following published protocol (41).

### Northern blot

RNA samples for northern blot were obtained after TRIzol extraction from cells infected at MOI 2 or higher and incubated for 48 hours (mammalian) or 72 hours (insect). Viral genome copies were quantified by RT-qPCR using a commercial kit (Primerdesign, ZIKV-R01042) and results were used to equalize sample loading into urea-PAGE. RNA fragments corresponding to the last 412, 328 and 240 nucleotides of the ZIKV genome were used as a reference size for sfRNA1, 2 and 3 and run together with samples. Transfer to nylon membrane was performed at 2 mA/1cm^2^ gel for 60 minutes. The rest of the procedure was based on commercial protocol available online (LI-COR Biotech, doc# 988-09394). Briefly, after the transfer, the membrane was UV-crosslinked and incubated overnight at 56°C with Biotin-16-UTP-labelled probe (UCCUCUAACCACUAGUCCCUCUUCUGGAGAUCCAC). After probe hybridization, the membrane was washed in low-(2x SSC with 0.1% SDS) and high-stringency buffers (0.5x SSC with 0.1% SDS), blocked (Odyssey Blocking Buffer, LI-COR Biotech, cat#927-70001), and incubated with Streptavidin-IRDye 800CW (LI-COR Biotech, cat#926-32230) for 30 min at room temperature. After washes the membrane was scanned with Odyssey CLx Imager (LI-COR Biotech).

### Expression and purification of recombinant Xrn1

Sequences for expressing human XRNI (amino acids 1-1173) (NP_061874.3) or Aedes aegypti XRNI (amino acids 1-1158) (XM_021853967.1) with an N-terminal 6xHis tag were cloned into pET29b(+) (Twist Biosciences) and transduced into Rosetta 2 (DE3) pLysS competent cells (Novagen). At OD600 cells were induced by addition of 0.5 mM IPTG and incubated for 16 hours overnight at 20 °C. Cells were harvested and lysed by sonication in 20 mM Tris pH8.5, 500 mM NaCl, 5 mM imidazole, 0.5% Triton X-100 and a protease inhibitor cocktail (Millipore). Protein was purified on NiNTA affinity resin (Cube Biotech) followed by ion exchange on a monoQ column (Cytiva) before being dialysis against 20 mM HEPES pH 7.5, 150 mM KCl and 1 mM TECP and concentrated to 5 mg/mL for storage at -70 °C.

### Xrn1 in *in vitro* digestion assay

Fragments 1 and 2 from ZIKV genome, for testing xrRNA1 and xrRNA2 respectively, were produced by PCR (Fragment 1 forward: <u>TAATACGACTCACTATAGG</u>CTGTGCCAGTTGACTGGGTTCCAACTGGG; Fragment 2 forward: <u>TAATACGACTCACTATAGG</u> GCACCAATCTTAATGT<u>CC</u>TCAGGCCTGCTAGTCAGCCAC; reverse: AGACCCATGGATTTCCCCACACCGGC) from the reverse genetic plasmids (WT_(AG)_ and PK2_mut(CC)_) and transcribed *in vitro* with in house-produced enzyme. The RNA fragments were incubated with Rpph (New England Biolabs, cat#M0356) following the manufacturers protocol for 45 minutes at 37°C and further incubated for 45 minutes at 37°C with equal amounts (1 μM) of recombinant Xrn1 from yeast (New England Biolabs, cat#M0338), mosquito or human (see section “Expression and purification of recombinant Xrn1”) using the NEB XrnI buffer. Product of digestions and sfRNA size controls (se section “Northern blot”) were resolved in urea-PAGE for visualization with gel documentation system.

